# Increased ventromedial prefrontal cortex activity in adolescence benefits prosocial reinforcement learning

**DOI:** 10.1101/2021.01.21.427660

**Authors:** Bianca Westhoff, Neeltje E. Blankenstein, Elisabeth Schreuders, Eveline A. Crone, Anna C. K. van Duijvenvoorde

## Abstract

Learning which of our behaviors benefit others contributes to social bonding and being liked by others. An important period for the development of (pro)social behavior is adolescence, in which peers become more salient and relationships intensify. It is, however, unknown how learning to benefit others develops across adolescence and what the underlying cognitive and neural mechanisms are. In this functional neuroimaging study, we assessed learning for self and others (i.e., prosocial learning) and the concurring neural tracking of prediction errors across adolescence (ages 9-21, N=74). Participants performed a two-choice probabilistic reinforcement learning task in which outcomes resulted in monetary consequences for themselves, an unknown other, or no one. Participants from all ages were able to learn for themselves and others, but learning for others showed a more protracted developmental trajectory. Prediction errors for self were observed in the ventral striatum and showed no age-related differences. However, prediction error coding for others was specifically observed in the ventromedial prefrontal cortex and showed age-related increases. These results reveal insights into the computational mechanisms of learning for others across adolescence, and highlight that learning for self and others show different age-related patterns.

## Introduction

Adolescence is a developmental phase that is characterized by transitions in social connections, and moreover, a phase during which social cognitive skills are acquired and/or improved (Blakemore & Mills, 2014; Casey et al., 2008; Crone & Dahl, 2012; Sawyer et al., 2018). As social acceptance and approval from peers often result from displaying prosocial behaviors, for adolescents establishing their social network it is key that they learn to help or benefit others (Steinberg & Morris, 2001). That is, to be able to behave in a prosocial manner, individuals need to learn which actions would result in positive outcomes for others. This type of learning is also referred to as prosocial learning (Lockwood et al., 2016; Sul et al., 2015). Generally speaking, learning from actions and outcomes is an important part of cognitive development and continues to improve in adolescence (Bolenz et al., 2017; Nussenbaum & Hartley, 2019; Peters, Braams, et al., 2014; Peters et al., 2016; van Duijvenvoorde et al., under revision). For adolescents, an especially salient environment that requires learning about the consequences of their actions is the interpersonal context (Blakemore & Mills, 2014; Nelson et al., 2005; Sawyer et al., 2018). Therefore, it is expected that especially prosocial learning shows improvements in adolescence. The goal of the current study was to unravel age-related differences in learning to benefit others using a prosocial learning context across adolescence.

The vast majority of recent neuroscientific studies investigating learning make use of formal reinforcement learning (RL) models. These models calculate individuals’ prediction errors (PEs) – the difference between expected and actual outcomes - over the course of learning. These PEs drive learning via a learning rate, which quantifies to what extent these PEs affect subsequent actions. Consequently, RL models and the resulting PEs enable studies to examine the neural tracking of value-guided decision making. Neuroscientific studies demonstrated that PE coding in a probabilistic reinforcement task context is associated with activation in the ventral striatum as well as the medial prefrontal cortex (see for reviews e.g., Cheong et al., 2017; Lockwood & Klein-Flügge, 2020; Olsson et al., 2020; Ruff & Fehr, 2014). Developmental studies using RL models found that adolescents show similar neural tracking of PEs as adults when learning stimulus-outcome associations, although the developmental patterns are inconsistent: some studies have reported elevated or lowered PE activity in the ventral striatum and connected structures in mid-adolescents relative to children and adults (Cohen et al., 2010; Davidow et al., 2016; Hauser et al., 2015; Jones et al., 2014), but this is not replicated in all studies (Christakou et al., 2013; van den Bos et al., 2012). Furthermore, age-related differences have been found in functional connectivity between the ventral striatum and ventral medial prefrontal cortex in relation to learning (van den Bos et al., 2012), suggesting that age-related improvements in learning are associated with stronger neural coupling between subcortical and cortical brain regions (van Duijvenvoorde et al., 2016, 2019). Taken together, previous studies point to the ventral striatum and medial prefrontal cortex as important brain areas for learning in non-social environments.

Previous studies investigating the neurocomputational mechanisms of prosocial learning have investigated whether the same neural signaling occurs for PEs for others as for self. Recently, in adults, it was found that PE tracking for both learning for others as for self occurred in the ventral striatum (Lockwood et al., 2016). However, PEs when learning for others were solely coded in the subgenual anterior cingulate cortex (sgACC), and these prosocial learning signals were predicted by cognitive empathy. That is, more empathic people showed more activity in de sgACC when learning to benefit others. Cognitive empathy – the ability to understand the emotional states of others (Netten et al., 2015; Pouw et al., 2013) - shows pronounced changes in adolescent development and relates positively to prosocial behaviors such as trust and reciprocity (Dumontheil et al., 2010; Eisenberg et al., 1995; van de Groep, Meuwese, et al., 2020). Therefore, we aimed to extend prior work by Lockwood and colleagues (2016) by investigating the neural tracking of PEs for others, and its relation with individual differences in cognitive empathy, in a sample with participants aged between 9 and 21 years.

In the current study, we adopted a prosocial learning task (Lockwood et al., 2016) in which participants could learn to obtain rewards for themselves, others, or no one. We administered this task to 74 adolescents between ages 9-21 years to examine age-related differences in learning for self and others, combined with functional neuroimaging (fMRI) for neural tracking of PEs. Based on prior studies, we performed regions-of-interest analyses for the ventral striatum, sgACC, and ventromedial prefrontal cortex (vmPFC). We expected that adolescents, similar to adults, would show PE related neural activity when learning both for self and others in the ventral striatum (Lockwood et al., 2016) and in the sgACC and possibly vmPFC when learning for others specifically (Christopoulos & King-Casas, 2015; Lockwood et al., 2016). For learning for self, research has remained inconclusive whether this activity peaks in mid-adolescence (Cohen et al., 2010; Davidow et al., 2016) or shows no age-related differences (van den Bos et al., 2012). Therefore we explored linear as well as non-linear (quadratic) age effects. We predicted that sgACC and vmPFC activity for prosocial learning would increase with age, based on prior studies showing age-related improvements in social-cognitive perspective-taking (Dumontheil et al., 2010). Finally, consistent with (Lockwood et al., 2016) we expected that individual differences in cognitive empathy would relate to neural tracking of PEs for others.

## Methods and Materials

### Participants

A total of 76 participants between ages 9 and 21 took part in this study. Participants were recruited through schools and local advertisements, as well as from participation in a previous study. Two participants were excluded from analyses because they were either diagnosed with a psychiatric disorder at the time of testing (*n*=1) or because the session was stopped due to discomfort in the scanner (*n*=1). We did not exclude participants based on task performance; there were no significant outliers on task performance (i.e., >3 SD) in any of the conditions. The final sample included 74 healthy participants (39 female, *M_age_* = 15.64, *SD_age_* = 4.18, range = 9.03 – 21.77 years, see Figure S1 for an overview of the number of participants across ages). The IQ scores, estimated with the Similarities and Block Design subtests of the WISC-III and WAIS-III, fell within the normal range (*M*_IQ_ = 110.24, *SD*_IQ_ = 10.37, range = 87.50 - 135.00), and did not correlate with age (*r*(72) = −0.11, *p* = .353).

The local institutional review board approved this study (reference: NL56438.058.16). Adult participants and parents of minors provided written informed consent, and minors provided written assent. All anatomical scans were cleared by a radiologist and no abnormalities were reported. Participants were screened for MRI contraindications and psychiatric or neurological disorders, and had normal or corrected-to-normal vision.

### Prosocial learning task

Participants played a two-choice probabilistic reinforcement learning task (prosocial learning task) in the MRI scanner (see Figure 1A). Participants were instructed to make a series of decisions between two pictures. One picture was associated with a high probability of winning 1 point, the other picture with a high probability of losing 1 point. The exact probabilities were 75% and 25% but were unknown to the participant. After the decision, participants were presented with the outcome to enable them to learn the reward contingencies.

**Figure 1.**
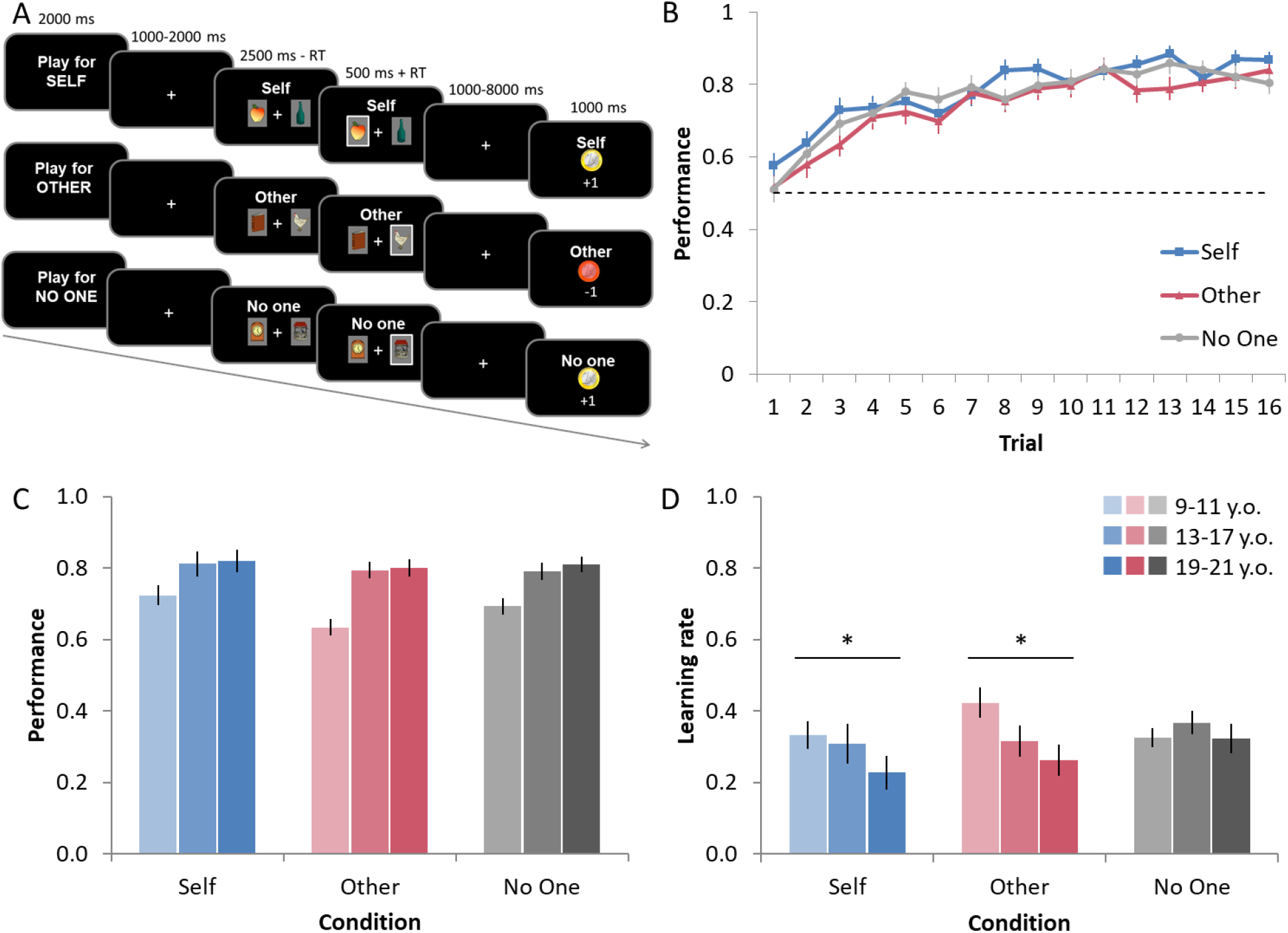
Prosocial learning task and behavioral data. **(A)** Participants played a two-choice probabilistic reinforcement learning task in which outcomes resulted in monetary consequences for themselves (Self condition), for an unknown other participant in the experiment who could not reciprocate (Other condition), or for No One. **(B)** Group-level performance across trials (learning curves) per condition, averaged across blocks. Performance represents the fraction of selecting the stimulus with a high reward contingency. The dashed line indicates performance at chance level (0.5). **(C)** Performance per condition per age cohort. In all conditions, performance improved across trials, but an age-related increase was only observed when learning for others. Note that age is used as a continuous variable in all analyses but is visualized as age cohorts for illustrative purposes. **(D)** Learning rates per condition per age cohort. Age-related decreased in learning rates are only observed in the Self and Other condition. Asterisks indicate significant effects of age. Error bars represent standard error of the mean (s.e.m.).

The participants played the task in three different conditions: for themselves (Self), for an unknown other participant (Other), or for No One. The latter condition was added as a control condition based on Lockwood et al. (2016). Participants were told that the other person was a peer also participating in the experiment who i) would not play the same game for them, ii) did not know who played for them (see Participant instructions in the Supplementary information). Each block started with an instruction screen that indicated who would receive the outcomes (Self, Other, or No One) for 2000 ms. This was followed by the presentation of two stimuli for 2500 ms during which participants were required to select one of these. The stimuli were common objects, such as chairs, apples, and shoes (see also (van den Bos, 2009)). If no response was given within the time frame, the text “Too late” appeared in the middle of the screen and these trials were excluded from analyses.

A selection frame around the chosen picture confirmed the response and remained visible for the duration of the interval and an additional 500 ms. A fixation screen (duration randomly jittered between 1000-2000 ms) preceded the outcome of their choice (+1 point or −1 point; 1000 ms). A randomly jittered fixation screen (1000-8000 ms) was shown after the outcome before the two pictures were presented again. The screen position of the stimulus (left or right) was counterbalanced across trials. Participants were instructed that the position of the stimulus did not matter, to encourage them to learn the reward contingencies regardless of stimulus position.

There were 144 trials in total, 48 for self, 48 for other, and 48 for no one, presented in three blocks of 16 trials. Each block began with a new pair of pictures. Participants completed three separate fMRI runs with a short break in between, each with one block of 16 trials per condition. The order of the conditions was counterbalanced across runs and between participants.

Participants were instructed that the total number of points in the Self condition was converted to money (each point valued €0.25), which they would get paid out on top of their flat participation rate (€20 for 9-11 y.o., €25 for 13-17 y.o., and €30 for 19-21 y.o.). The minimum of this extra amount of money was €1 to avoid null scores, and the maximum was €12. Additionally, participants were instructed that their choices in the Other condition were paid out to a participant entering the experiment after them. Consequently, participants received an additional fee from a participant before them in the experiment (minimum €1 and a maximum €12), but only at the end of the experiment. Finally, it was instructed that choices in the No One condition had no financial consequences.

### Questionnaires

To assess cognitive empathy, participants completed the Interpersonal Reactivity Index (IRI; (Davis, 1983)). This widely used self-report questionnaire consists of 4 subscales (Perspective-Taking and Fantasy as cognitive empathy subscales; and Personal Distress and Empathic Concern as affective empathy subscales) with 6 items each. To create a measure of cognitive empathy, two subscales were combined (Pulos et al., 2004): the Perspective-Taking subscale (e.g., “I sometimes try to understand my friends better by imagining how things look from their perspective”, Cronbach’s alpha = 0.710) and the Fantasy subscale (e.g., “I really get involved with the feelings of the characters in a novel”, Cronbach’s alpha = 0.786). All items can be answered on a five-point Likert scale ranging from (0) not true at all to (4) completely true, and higher scores indicate higher levels of empathy. We used a Dutch adolescent version for all ages in our study, with items adapted for the youngest ages in the study (Hawk et al., 2013).

### Procedure

Participants were accustomed to the MRI environment using a mock scanner, and received instructions on the prosocial learning task in a quiet laboratory room. Instructions for the task were displayed on a screen and read out loud by an experimenter. Participants completed 8 practice trials in each condition. In the scanner, participants responded with their right hand using a button box. Head movements were restricted with foam padding. The fMRI scan was accompanied by a high-definition structural scan. Questionnaires were filled out at their home prior to the scanning session, via Qualtrics (www.qualtrics.com).

### Computational modeling of behavioral data

#### Model fitting

First, we obtained PEs and learning rates from the behavioral choice data, which were subsequently used in behavioral and fMRI analyses. We modeled learning behavior in the Self, Other, and No One conditions separately, using a standard reinforcement learning model (similar to Lockwood et al., 2016). Simple RL models state that the expected value of a future action (Q_t+1_(i)) should be a function of current expectations (Q_t_(i)) and the difference between the actual reward that has been experienced on this trial (R_t_). The learning rate α, bounded between 0 and 1, determines how much the value of the chosen stimulus is updated based on the new outcome. In particular, the learning rate parameter speeds up or slows down the acquisition and updating of associations. Given the stable probabilistic associations between stimulus and outcome, lower learning rates may result in more optimal learning than very high learning rates (Davidow et al., 2016).

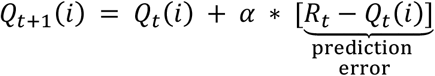

To select an action based on the computed values, we used a standard softmax choice function. For a given set of parameters, this equation allows us to compute the probability of the next choice being “i”:

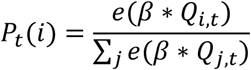

Beta (β) is modeled as an inverse temperature parameter and determines *how strongly* action probabilities are guided by their expected values (Q). With larger β, actions are more deterministic and driven by expected values, resulting in selecting the option with the highest value. With a lower β, actions are more random or exploratory. This inverse temperature parameter β thus affects errors, where a decrease will lead to more random (i.e., less driven by expected values) choices. β did not differ between conditions, although with age people were more strongly driven by expected values (see Figure S2 for the β across age cohorts for each condition).

We used the maximum a posteriori (MAP) approach (Daw, 2011) for fitting the RL model to participants’ choices per condition. To facilitate stable estimation across subjects, we used weakly informative priors to regularize the estimated priors toward realistic ones. These weakly informative priors were based on previous research (den Ouden et al., 2013), and included a Beta distribution for the estimated α parameter and a Gaussian distribution for the estimated β parameter.

#### Model comparison

Based on previous developmental findings (e.g.,(van den Bos et al., 2012)) we compared an alternative model with two learning parameters (i.e., separate learning rates for gains and losses) in order to benchmark the performance of the one-learning parameter model (i.e., one learning rate). Model comparisons revealed that the one-learning parameter model had a superior fit to the behavioral data for each condition, according to the Bayesian Information Criterion (BIC) (see Figure S3). This was the case for the majority of the participants (82.4% Self, 74.3% Other, 76.7% No One), in all age cohorts.

#### Simulations and parameter recovery

To assess whether computational model parameters could be successfully recovered, we simulated choice behavior for a range of learning rates. Parameter recovery is acceptable and is shown in Figure S4. See supplemental information for a more detailed description of the used procedures.

### Behavioral analyses

To assess learning for Self, Other, and No One, and their developmental patterns in the prosocial learning task, we fitted logistic generalized linear mixed models (GLMMs) to decisions (correct coded 1, incorrect 0) for each condition separately. These analyses were conducted in R version 4.0.1 (R Core Team. 2020), using the lme4 package (Bates et al., 2014). Our GLMMs included fixed effects of Age in years (linear and quadratic), Condition, Trial, and all interactions. In all models, participant ID entered the regression as a random effect to handle the repeated nature of the data. Where applicable, Trial and Condition were additionally included as random slopes per subject. We performed post hoc tests per condition to delineate the Age x Trial x Condition effects. In all models, continuous independent variables are mean-centered and scaled, and categorical predictor variables were specified by a sum-to-zero contrast (e.g., sex: −1 = boy, 1 = girl). For the mixed-effects model analyses, the optimizer “bobyqa” was used (Powell, 2009) with a maximum number of 1×10^5^ iterations. P-values for all individual terms were determined by Loglikelihood Ratio Tests as implemented in the mixed function in the afex package (Singmann et al., 2020).

Moreover, we examined how people updated the value of stimuli based on outcomes for Self, Others, and No One. Therefore, we tested differences between conditions in learning rates with repeated measures ANCOVA’s with three levels (Self, Other, No One) and linear age as a covariate. In cases in which the assumption of sphericity was violated, Greenhouse-Geisser corrections were reported. Finally, we assessed whether cognitive empathy related to learning performance, learning rate, and PE activation when learning for Others. We ran separate robust linear regression analyses (5000 bootstraps), each with age linear and cognitive empathy as predictors. The latter analyses were performed in SPSS 25 (IBM Corp.), and all tests are two-sided.

### fMRI acquisition

For acquiring (functional) MRI data, we used a 3T Philips scanner (Philips Achieva TX) with a standard eight-channel whole-head coil. The learning task was projected on a screen that was viewed through a mirror on the head coil. Functional scans were acquired during three runs of 200 dynamics each, using T2* echo-planar imaging (EPI). The volumes covered the entire brain (repetition time (TR) = 2.2 s; echo time (TE) = 30 ms; sequential acquisition, 38 slices; voxel size 2.75 × 2.75 × 2.75 mm; field of view (FOV) = 220 (ap) x 220 (rl) x 114.68 (fh) mm). The first two volumes were discarded to allow for equilibration of T1 saturation effects. After the learning task, a high-resolution 3D T1 scan for anatomical reference was obtained (TR = 9.76 msec, TE = 4.95 msec, 140 slices, voxel size = 0.875 × 0.875 × 0.875 mm, FOV = 224 (ap) x 177 (rl) x 168 (fh) mm).

#### Preprocessing

Data were analyzed using SPM8 (Wellcome Department of Cognitive Neurology, London). Images were corrected for slice timing acquisition and rigid body motion. We spatially normalized functional volumes to T1 templates. Occasional framewise displacement >3mm occurred for 3 participants in 1-2 volumes. For those participants with frame-frame head motion >3mm, an extra regressor was included corresponding to each volume (*n* = 3, for maximum 2 volumes). All other participants did not exceed translational head movement more than 3mm in any of the scans (*Mean* = 0.65mm, *SD* = 0.059mm). The normalization algorithm used a 12 parameter affine transform with a nonlinear transformation involving cosine basis function, and resampled the volumes to 3 mm^3^ voxels. Templates were based on MNI305 stereotaxic space. The functional volumes were spatially smoothed using a 6 mm full width at half maximum (FWHM) isotropic Gaussian kernel.

#### General-linear model

We used the general linear model (GLM) in SPM8 to perform statistical analyses on individual subjects’ data. The fMRI time series were modeled as a series of two events: the decision phase (Expected Value, EV) and the outcome phase (PE), convolved with a canonical hemodynamic response function (HRF). The onset of the choice (EV), and the onset of the outcome (PE) were both modeled with zero duration. Each of these regressors was associated with a parametric modulator taken from the computational model. At the time a stimulus was selected (decision phase) this was the chosen expected value, and at the time of the outcome, the PE. The PEs were estimated using each subject’s own alpha and beta from each condition. Trials on which participants did not respond were modeled separately as a regressor of no interest. Six motion parameters, and -if applicable-motion censoring regressors were included as nuisance regressors. We used the MarsBaR toolbox (Brett, Anton, Valabregue, & Poline, 2002; http://marsbar.sourceforge.net) to visualize the patterns of activation, in clusters identified in the whole-brain results. Coordinates of local maxima are reported in MNI space. Our main hypotheses centered on PE coding. For condition effects, we examined contrasts of Self versus Other in concordance with (Lockwood et al.(2016). Contrasts were obtained from a flexible factorial design with three levels (Self PE, Other PE, No One PE). Effects and conclusion remained the same when testing Self PE > Other PE + No One PE, and Other PE > Self PE + No One PE (see Supplemental Table S2). Age effects (linear and quadratic) were tested in follow-up regressions.

#### ROI selection and fMRI analyses

The a priori regions of interest (ROI) in which we test our main hypotheses were defined anatomically and based on previous research on (prosocial) learning (Lockwood et al., 2016; van den Bos et al., 2012; van Duijvenvoorde et al., 2014). In concordance with previous studies, masks were taken from an appropriate atlas. That is, the bilateral ventral striatum and vmPFC were determined by an anatomical mask from the Harvard-Oxford Atlas (Braams et al., 2015; Peters & Crone, 2017), and the sgACC was defined as Brodmann areas (BA) 25 and s24 (Lockwood et al., 2016). The sgACC region and the ventral striatum are anatomically adjacent and partly overlapping (see Figure S5), but significant peak activations in either ROI were not observed in these overlapping voxels. Coordinates for local maxima are reported in MNI space. Effects in our ROIs are reported at *p* < .05 FWE-small volume corrected (SVC). Predictions were tested while correcting for multiple comparisons (3 ROIs) by limiting the false discovery rate (FDR; (Benjamini & Hochberg, 1995)); all reported tests survived this correction. Explorative whole-brain analyses are reported in Supplemental Tables S1 and S2 with a *p* < .05 FWE correction on voxel level (Figure S6).

## Results

### Developmental differences in learning to obtain rewards for Self, Others, or No One

Results showed at the group level, participants were able to learn for Self, Other, and No One, as they performed above chance level in all conditions (0.5; *t* values > 13.0, *P* values < 0.001, *df* = 73; Figure 1B). Using a generalized linear mixed model (GLMM) on participants’ choice behavior over trials, we assessed age-related differences in performance when learning for Self, Other, and No One. A main effect of age showed that performance in the learning task improved linearly with age (main effect of Age linear, B = 0.38, *p* < .001), there were no quadratic age effects. Moreover, we observed that age-related differences in learning curves differed per condition (Age linear x Condition x Trial, *p* = .006). Post-hoc analyses per condition revealed that in all conditions, performance improved across trials (main effect of Trial, all *P*s < 0.001), but an age-related increase was only observed when learning for others (Age linear x Trial, for Other condition, B = 0.266, *p* < .001; Self and No One conditions: *P*s > 0.2; Figure 1C and Figure S7. Together, these findings suggest that across adolescence prosocial learning shows a more protracted improvement than when learning for self or no one.

Next, we examined how people updated the value of stimuli on the basis of outcomes for Self, Others, and No One. That is, higher learning rates indicate that people adjusted behavior quickly towards recent feedback, whereas lower learning rates indicate a slower pace in updating, integrating outcomes across multiple trials. Overall, learning rates decreased linearly with age (main effect of Age linear, *F*(1,72) = 5.99, *p* = .017, *η*_p_^2^ = 0.077). Even though learning rate parameters did not differ between conditions (main effect of Condition, *F*(1.74, 125.16) = 2.90, *p* = .069, *η*_p_^2^ = 0.039), there was a significant age x condition interaction (Age linear x Condition, *F*(1.738, 125.16) = 3.33, *p* = .045, *η*_p_^2^ = 0.044). Specifically, the age-related decreases in learning rates were only observed in the Self and Other conditions (Figure 1D), but not in the No One condition. These findings show that for both learning for self and others, younger participants responded more to recent feedback, whereas older participants integrated feedback more over trials.

### Identifying common and distinct coding of prediction errors for Self and Others

To formally investigate the brain regions that were responding to PEs for Self, Others, and No One, we conducted a conjunction analysis to explore whether there were regions that commonly code PEs across all conditions. Common activation for PEs regardless of the beneficiary was observed in the vmPFC (MNI coordinates [*x* = −9, *y* = 44, *z* = −11], *Z* = 5.33, *k* = 136, *p* < .001, SVC-FWE), ventral striatum ([*x* = −9, *y* = 11, *z* = −11], *Z* = 5.05, *k* = 23, *p* < .001, SVC-FWE, and [*x* = 12, *y* = 14, *z* = −8], *Z* = 4.43, *k* = 18, *p* < .001, SVC-FWE), and sgACC ([*x* = −6, *y* = 14, *z* = −8], *Z* = 5.47, *k* = 32, *p* < .001; and ([*x* = 6, *y* = 17, *z* = −8, *Z* = 4.60, *k* = 21, *p* = .001; and ([*x* = 9, *y* = 8, *z* = −14, *Z* = 3.67, *k* = 2, *p* = .029, SVC-FWE) (Figure 2). These findings show that all regions of interest were involved in PE coding, in each condition.

**Figure 2.**
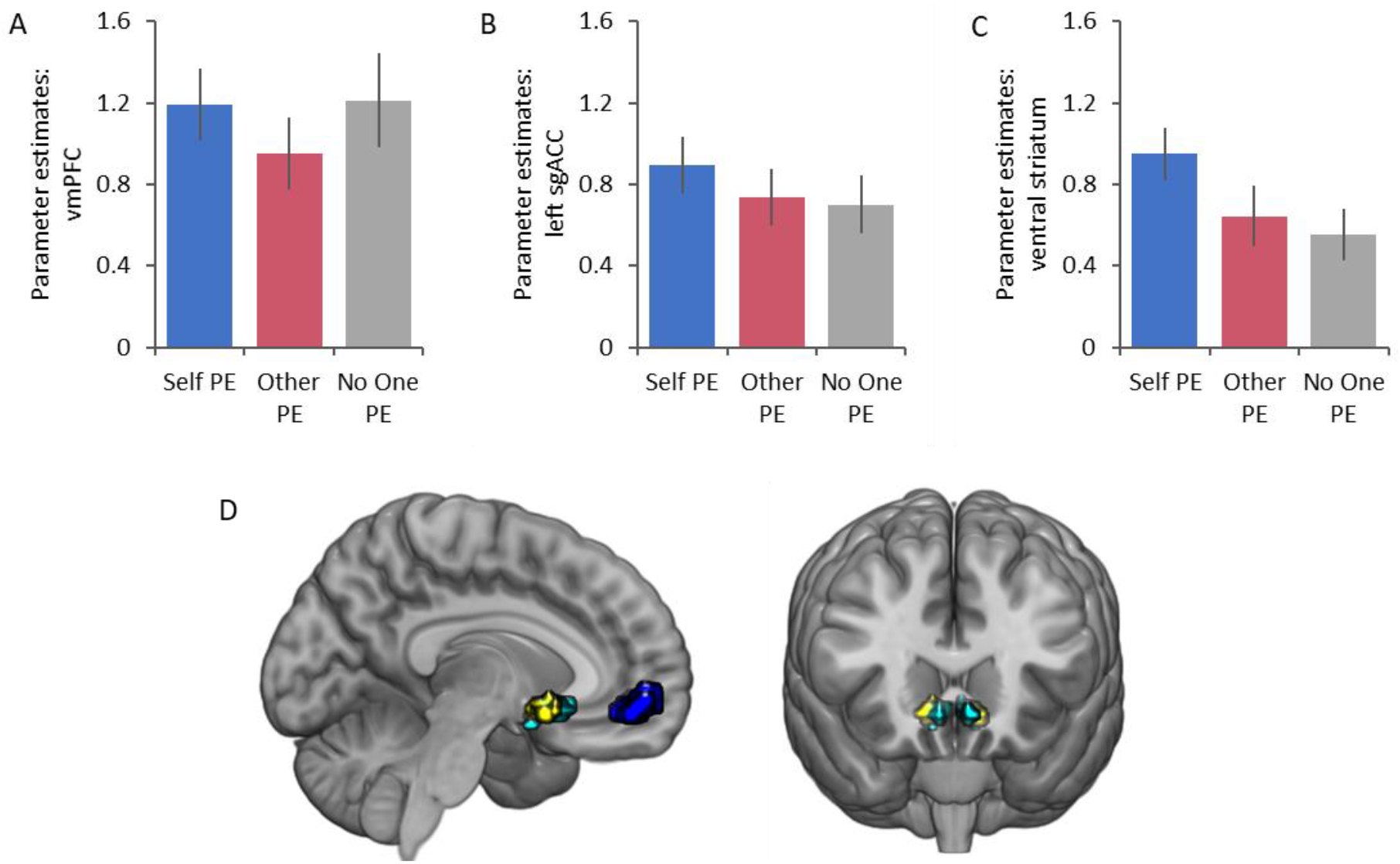
Common prediction error (PE) coding in three regions of interest. Shown are the responses to prediction errors for Self, Other, and No One in **(A)** the vmPFC, **(B)** left sgACC, and **(C)** ventral striatum. **(D)** Significant clusters of activation in the vmPFC (blue), sgACC (cyan), and ventral striatum (yellow). All images displayed at *p* < .05 FWE-SVC.

Next, we examined which brain regions responded more to PEs for Self than for Other by contrasting the Self condition against the Other condition (see supplemental information for contrasts including the No One condition). The left ventral striatum was the only region to respond more strongly to PEs for Self ([*x* = 12, *y* = 11, *z* = −11], *Z* = 4.37, *k* = 9, *p* < .001, SVC-FWE; Figure 3). When examining effects of age we observed no linear or quadratic age-related differences in self-related PE coding. These findings indicate that the ventral striatum responds more to PEs for Self than for Others, and this effect did not differ across adolescence.

**Figure 3.**
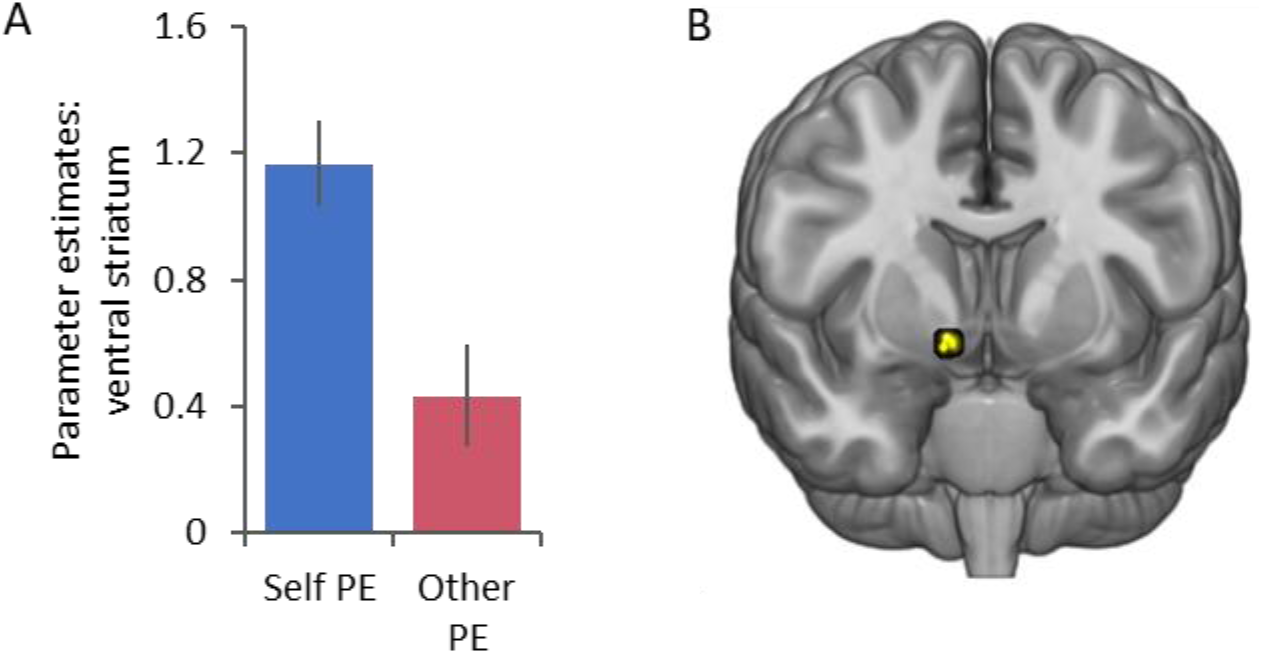
Ventral striatum response to prediction errors for Self versus Other. **(A)** Left ventral striatum [x=12, y=11, z=-11] response for Self PE and Other PE. **(B)** Overlay of the response for Self PE > Other PE in the left ventral striatum. All images displayed at *p* < .05 FWE-SVC.

We next identified regions that corresponded to PEs for others exclusively by contrasting the Other condition against the Self condition. No voxels in our ROIs responded more strongly to prosocial PEs. When adding age (linear and quadratic) to the model to examine whether age-differences were related to prosocial PE coding, we observed that with age, vmPFC increasingly responded to prosocial PEs ([*x* = −15, *y* = 50, *z* = 8], *Z* = 4.95, *k* = 45, *p* = .004, SVC-FWE; Figure 4). No effects of quadratic age were observed. This shows that the vmPFC is increasingly involved in prosocial PE coding across adolescence.

**Figure 4.**
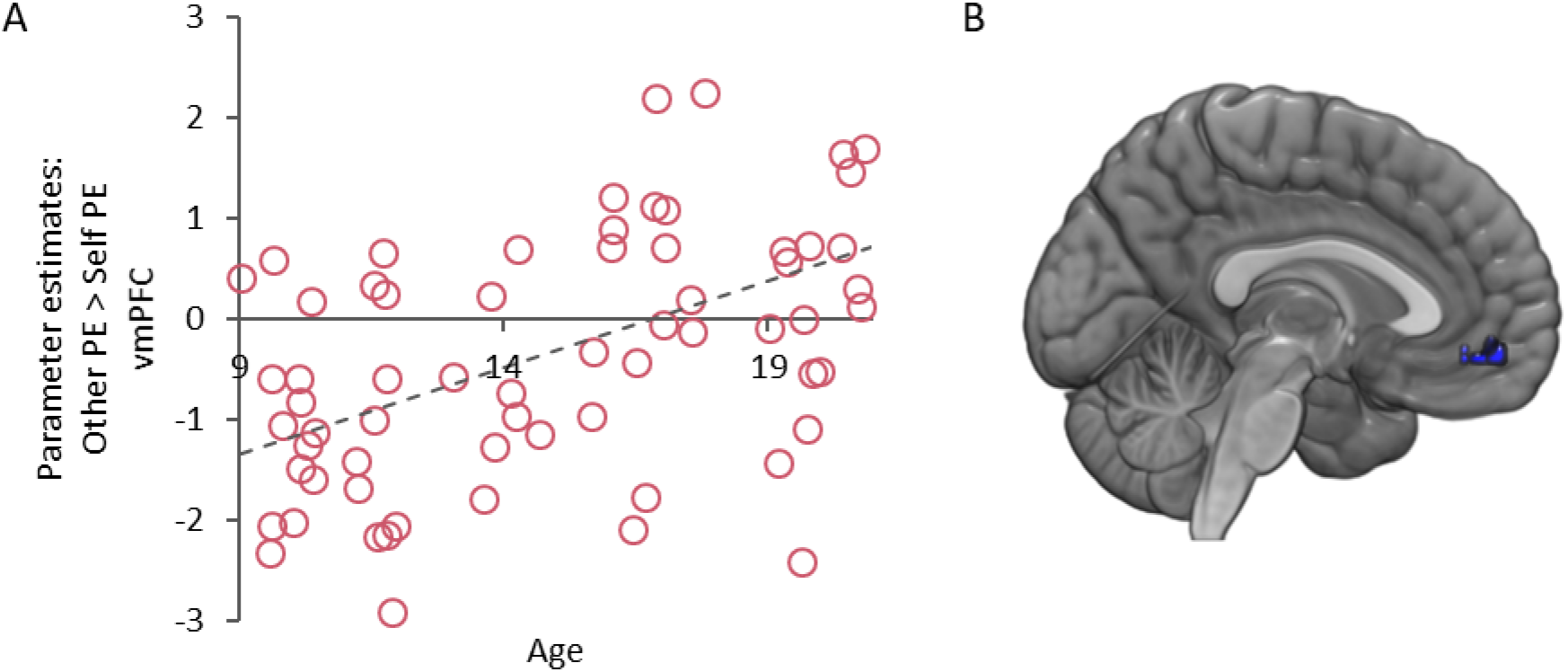
Linear age effects in responses to Other PE > Self PE in the vmPFC. **(A)** scatterplot showing the relation between age and activation in the vmPFC for Other PE > Self PE. **(B)** Overlay of the response for Other PE > Self PE in the vmPFC [−15, 41, −11]. All images displayed at *p* < .05 FWE-SVC.

### Links between cognitive empathy and learning for Others

Finally, we examined the link between cognitive empathy and prosocial learning using robust linear regression analyses (5000 bootstraps). First, we assessed whether cognitive empathy predicted performance for Other, while controlling for performance for Self and age (linear). We observed that individuals with higher empathy ratings, showed better prosocial learning performance (cognitive empathy, *b* = − 0.019, β = − 0.264, *P* = 0.019, 95% CI = [− 0.036, −0.003], see Figure 5A). Subsequently, we assessed whether cognitive empathy predicted learning rate in the Other condition (controlled for learning rate in the Self condition and age (linear)). Results showed that individuals with higher empathy ratings had lower learning rates when learning for Others (cognitive empathy, *b* = 0.011, β = 0.217, *P* = 0.016, 95% CI = [0.002, 0.020], see Figure 5B). Together, these findings indicate that individuals with more empathy show better learning performance, and integrate information more over trials when learning to benefit others. Finally, we assessed the relation between cognitive empathy and the prosocial PE coding in the vmPFC. For this purpose, we extracted the values of the Other PE > Self PE contrast in vmPFC that showed age-related change (Figure 4). Results showed that greater Other-related PE activation in the vmPFC related to higher empathy scores (cognitive empathy, *b* = 0.112, β = 0.047, *P* = 0.020, 95% CI = [0.018, 0.206]). However, when controlling for age, this relation was no longer significant (cognitive empathy, *b* = 0.046, β = 0.110, *P* = 0.291, 95% CI = [−0.040, 0.131]).

**Figure 5.**
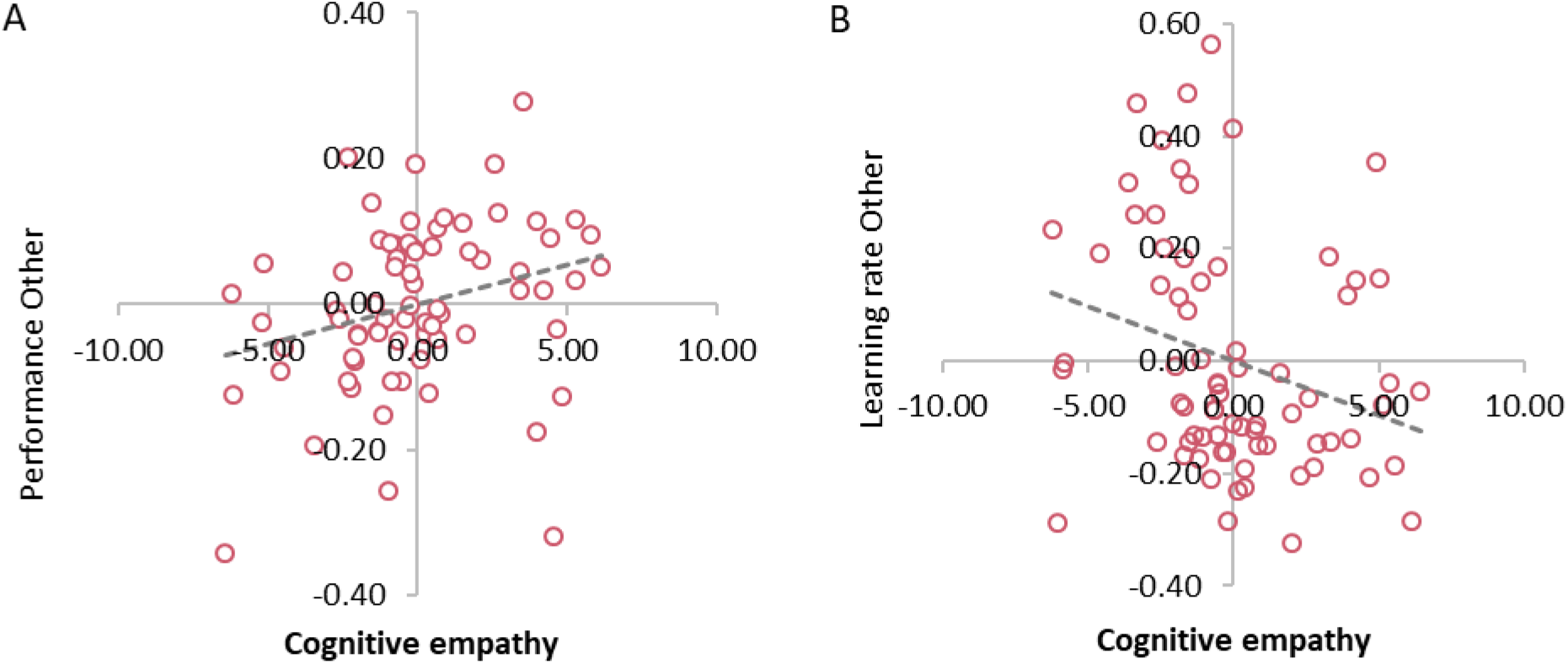
Relation of cognitive empathy with performance for Others and learning rate for Others. **(A)** Partial regression plot showing that individuals with more cognitive empathy perform better for Others (controlled for performance for Self and linear age). **(B)** Partial regression plot showing that individuals with more cognitive empathy have lower learning rates when learning for Others (controlled for learning rate for Self and linear age).

## Discussion

This study examined the developmental trajectories of prosocial learning and self-related learning in an adolescent sample spanning ages 9-21 years. We examined the underlying mechanisms in this developmental sample by assessing the neural tracking of PEs during learning for self and others, and how individual differences in cognitive empathy relate to prosocial learning performance. To this end, participants played a two-choice probabilistic reinforcement learning task in which outcomes resulted in monetary consequences for themselves (Self) or an unknown other (Other; prosocial). Our results show improvements in learning for self and others, but the developmental trajectory of prosocial learning is more protracted compared to learning for self. PEs for self were related to activation in the left ventral striatum, which did not show age-related differences. On the other hand, vmPFC-related PE activation during prosocial learning increased with age, and related to individual differences in cognitive empathy. Together, these findings highlight that learning for self and others show different age-related patterns.

The main goal of this study was to examine age-related differences in prosocial learning. Behaviorally, we observed that it is not until mid-adolescence that participants learn similarly for themselves and others. These findings may suggest a stronger self-bias in younger adolescents (van der Aar et al., 2018), and stronger motivation to learn for self and others in older ages (Apps et al., 2016). Neurally, we observe that a reward-related network including the ventral striatum, sgACC, and vmPFC respond significantly to PEs when learning for self, others, and no one. This conjunction presented the starting point for our interest in testing condition-specific learning effects. Contrary to Lockwood et al. (2016), who observed similar PE neural tracking values in the ventral striatum for learning for self and others in adults, we observed that PE neural tracking was stronger in the ventral striatum for self than for others. Recent reviews, however, suggest that the striatum is related to a range of computations that take place during social learning that could reflect both self-related and other-related learning (Joiner et al., 2017), or the difference between winning for self and others (Báez-Mendoza & Schultz, 2013). Therefore, one explanation for our findings could be related to the possible stronger self-focus or the greater focus on social comparisons reflected in the ventral striatum that may be specific to this younger population.

Learning for others, compared to learning for self, was associated with stronger activation in the vmPFC. Previous prosocial reinforcement learning studies have suggested that the vmPFC is, however, most responsive to self-related reward processing (Sul et al., 2015) and self-representation (Sui & Humphreys, 2017), or does not differentiate between self and other-related PEs (Lockwood et al., 2016). On the other hand, the vmPFC is suggested to respond to prosocial rewards in adults (Christopoulos & King-Casas, 2015), to others’ outcome PEs (Burke et al., 2010), and to simulated others’ reward PEs (Suzuki et al., 2012). These contradictory findings suggest that the encoding of PEs for others *and* self might be tightly linked in brain regions such as the ventral striatum and the vmPFC (Joiner et al., 2017). In this first developmental sample investigating prosocial learning, we observe a specificity for self in the ventral striatum and an increased specificity for prosocial PE coding in the vmPFC. Alternatively, the pattern of age-related differences we observed for other and self-learning in the vmPFC may also support the perspective of a decreasing self-focus with age. For instance, previous work on self-concept development highlights that perspectives of others and self become more merged across development (van der Cruijsen et al., 2019). However, longitudinal studies are more powerful and essential for examining the true developmental trajectories of prosocial learning.

Besides the age-related differences in other-related learning, we observed that consistent with previous findings (Lockwood et al., 2016), individual differences in cognitive empathy were related to prosocial learning. Individuals with higher levels of empathy performed better for others and integrated outcomes more over time. However, prosocial PE coding in the brain and empathy were not related. Possibly, age-related differences in brain activity during prosocial PE tracking are explained by other mechanisms. For instance, although there was no reciprocity or competition, participants may have been influenced by social inequality preferences, such as a dislike to getting more (i.e., advantageous inequality aversion), or less (i.e., disadvantageous inequality aversion) than the other participant (Dawes et al., 2007; Fehr & Schmidt, 1999; Meuwese et al., 2015; Westhoff et al., 2020). Future studies could more explicitly assess several social-cognitive skills, strategies, and motivations along with a prosocial learning task to examine what behavioral mechanisms rely most on adolescents’ prosocial learning.

Prior developmental studies on general reinforcement learning remained inconclusive about whether age-related differences were observed in PE neural tracking in the ventral striatum (Christakou et al., 2013; Cohen et al., 2010; Hauser et al., 2015; van den Bos et al., 2012). Here, age-related differences in PE coding for self were not observed in the ventral striatum. In contrast to other studies (Cohen et al., 2010; Peters & Crone, 2017) we also did not find any quadratic age effects in learning or PE coding. This is possibly due to our narrower age range (9-21 y.o. instead of 8-30 y.o.), as another developmental study on learning also has not observed age-related changes in ventral striatum activity in a similar age range (van den Bos et al., 2012). Indeed, a recent review recommended using samples with wider age ranges, including children and adults, when examining quadratic age effects across adolescence (Li, 2017). It should be noted, however, that although we did not find age-effects in the ventral striatum, the behavioral learning performance for self showed linear improvements with age. This could also indicate that other mechanisms than simple PE coding may be related to behavioral learning improvement over time within the current age range. For example, a prior study in young adults indicated that besides well-known model-free learning, another more sophisticated and flexible learning system is model-based learning. These two distinct computational strategies use different error signals which are computed in partially distinct brain areas (Gläscher et al., 2010). Moreover, it has been found that people may use different learning strategies, which show different neural activation patterns (Peters, Koolschijn, et al., 2014). Future studies are needed to assess whether age-related improvements in learning performance may be more strongly related to strategic learning differences.

The current study had several limitations that should be addressed in future research. First, prosocial learning was restricted to unknown others and should be extended in future research to other beneficiaries. Previous studies have shown that prosocial behaviors and their neural correlates in adolescence strongly depend on the beneficiary (e.g., (Brandner et al., 2020; Schreuders et al., 2018; van de Groep, Zanolie, et al., 2020; Westhoff et al., 2020). Whether such differences between beneficiaries are also visible in prosocial *learning* and the concurrent neural tracking of PEs, is an interesting question for future studies. Second, the neural results for the no one condition showed an intermediate pattern between learning for self and others, which is difficult to interpret. Behavioral analyses showed that participants generally performed well in this condition (i.e., not significantly different from learning for self), even though no monetary reinforcers were given depending on task performance. Although including the no one conditions in our contrasts of interest did not alter our main findings, this condition was possibly interpreted by participants in different ways, in which some participants were internally motivated to perform well (e.g., (Satterthwaite et al., 2012).

In conclusion, we found that prosocial learning showed age-related improvements across adolescence, suggesting a developmental shift from self-focus in early adolescence to self and other-focus in late adolescence and early adulthood (Crone & Fuligni, 2020). This developmental improvement was associated with stronger recruitment of the vmPFC. This study has implications for learning in social settings, such as educational contexts (Altikulaç et al., 2019), as well as for how children develop prosocial values when learning for unknown others. This study provides the first building blocks to understand age-related differences in how adolescents learn to benefit others.

## Acknowledgements

We would like to thank all participants and their parents for their cooperation. We thank Myrna van den Berg, Sander de Heer, and Iris Koele for all their efforts during data collection. We thank Kiki Zanolie and Jochem Spaans for help with programming the task, and Ili Ma for help with the model simulations and parameter recovery. This work was supported by the European Research Council (ERC) under the European Union’s Horizon 2020 research and innovation programme [grant number 681632, to E.A.C]; and by the Netherlands Organization for Scientific Research (NWO) [Open Research Area grant, grant number 464-15-176, to A.C.K.D]. The funder had no role in the conceptualization, design, data collection, analysis, decision to publish, or preparation of the manuscript.

## Author contributions

**B.W.**: Conceptualization; Data curation; Formal analysis; Investigation;; Project administration; Visualization; Writing - original draft; Writing - review & editing. **N.E.B.**: Investigation; Writing - review & editing. **E.S.**: Investigation; Writing - review & editing. **E.A.C.**: Conceptualization; Funding acquisition; Resources; Writing - original draft; Writing - review & editing. **A.C.K.D.**: Conceptualization; Data curation; Formal analysis; Funding acquisition; Resources; Software; Supervision; Writing - original draft; Writing - review & editing. All authors approved the final version of the manuscript.

