## Supplementary Information for "Increased ventromedial prefrontal cortex activity in adolescence benefits prosocial reinforcement learning"

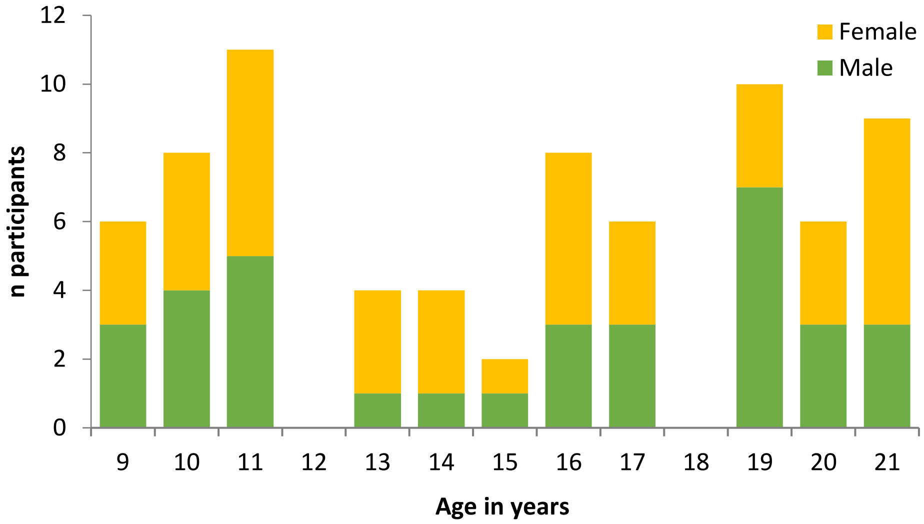


**Figure S1. Number of participants across age per sex.** In total, 74 participants were included (39 female, 35 male).

**Inverse temperature**

The inverse temperature parameter was examined as an index to what extent people followed expected value in their choice behavior, and is also considered a parameter of decision noise. Decision noise did not differ between conditions (main effect of Condition, F(1.83, 131.70) = 2.39, $p$ = .100, $\eta_p^2$ = 0.032). However, with increasing age, decision noise decreased linearly (main effect of Age, $F$(1, 72) = 11.24, $p$ = .001, $\eta_p^2$ = 0.135, see Figure S2), although this age effect did not differ per condition (Age x Condition, $F$(1.83, 131.70) = 1.995, $p$ = .144, $\eta_p^2$ = 0.027).

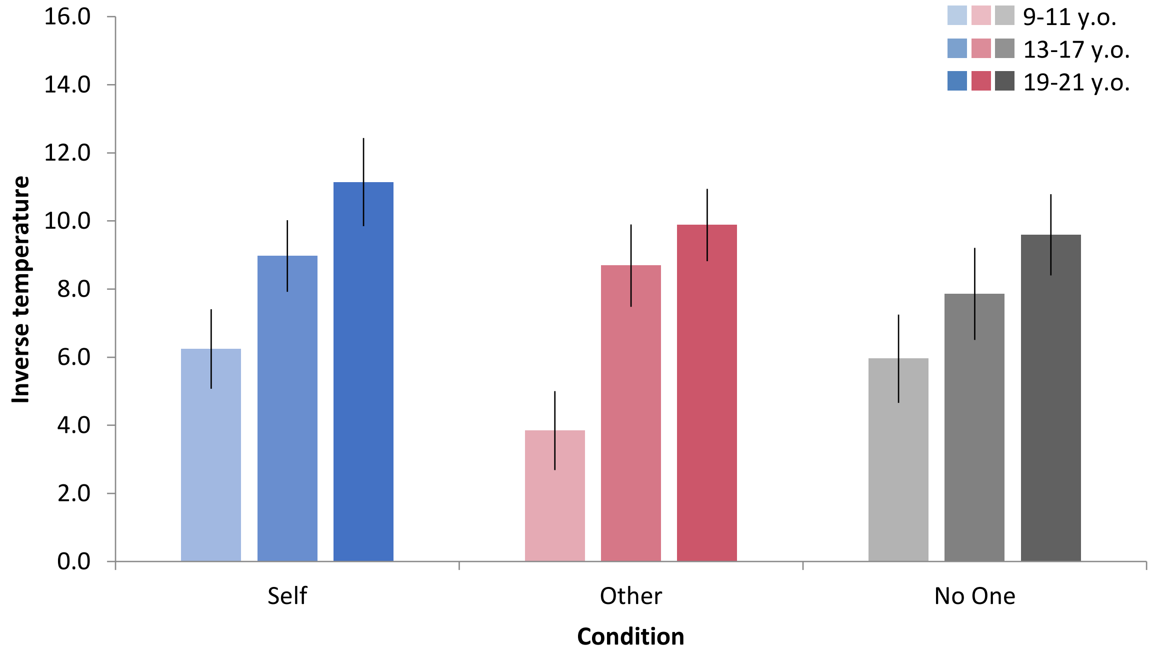


**Figure S2. Inverse temperature per condition per age cohort.** Age is used as a continuous variable in all analyses, but is visualized as age cohorts for illustrative purposes and interpretability.

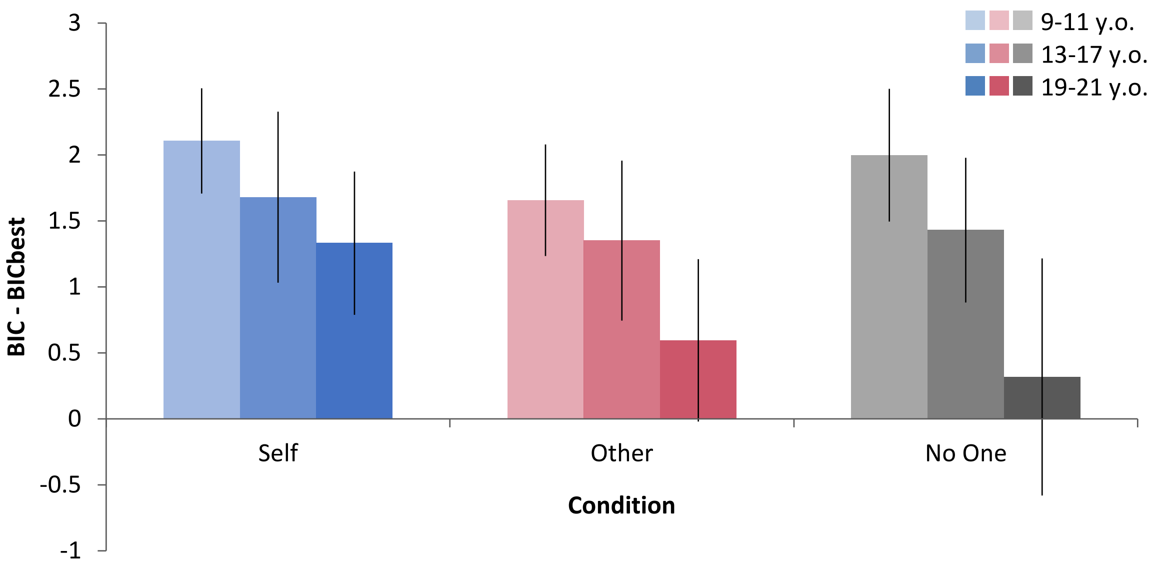


**Figure S3. BIC values per condition per age cohort.** Bars show BIC differences of the model with two learning rates (gain and loss) with the best model (one learning rate). BIC values were calculated per participant and were averaged per age cohort. Lower BIC values indicate better model fit. BIC values are shown separately per age cohort and per condition, showing that for all age cohorts and all conditions the best model has one learning rate. Error bars represent standard error of the mean.

### Simulations and parameter recovery

To test whether the learning rates could be recovered, we simulated choice behavior for a range of learning rates (i.e., α = 0.2, 0.4, 0.6, and 0.8) β values (i.e., 3, 6, and 9). For each combination of α and β, we simulated 100 subjects, using gaussian noise in these parameters. Next, on this simulated data, we estimated α and β following the MAP approach as described above (See *Computational modeling of behavioral data; Model fitting*). Parameter recovery is shown in Figure S4. The recovery of the learning rates seems fair (Figure S4-A – S4-C), with less extreme values showing better recovery. This pattern was observed for each combination of β values (i.e., 3, 6, 9).

Additionally, we assessed whether a simulation with values based on real participant data would also reveal a good recovery of parameters. As such, we simulated a new participant dataset based on the α and β values from our participants as input parameters. This resulted in a simulated dataset with 74 participants. Parameter recovery for the learning rates per condition is shown in Figure S4D. Together, these simulations suggest that our behavioral task and modelling procedure seem suitable.

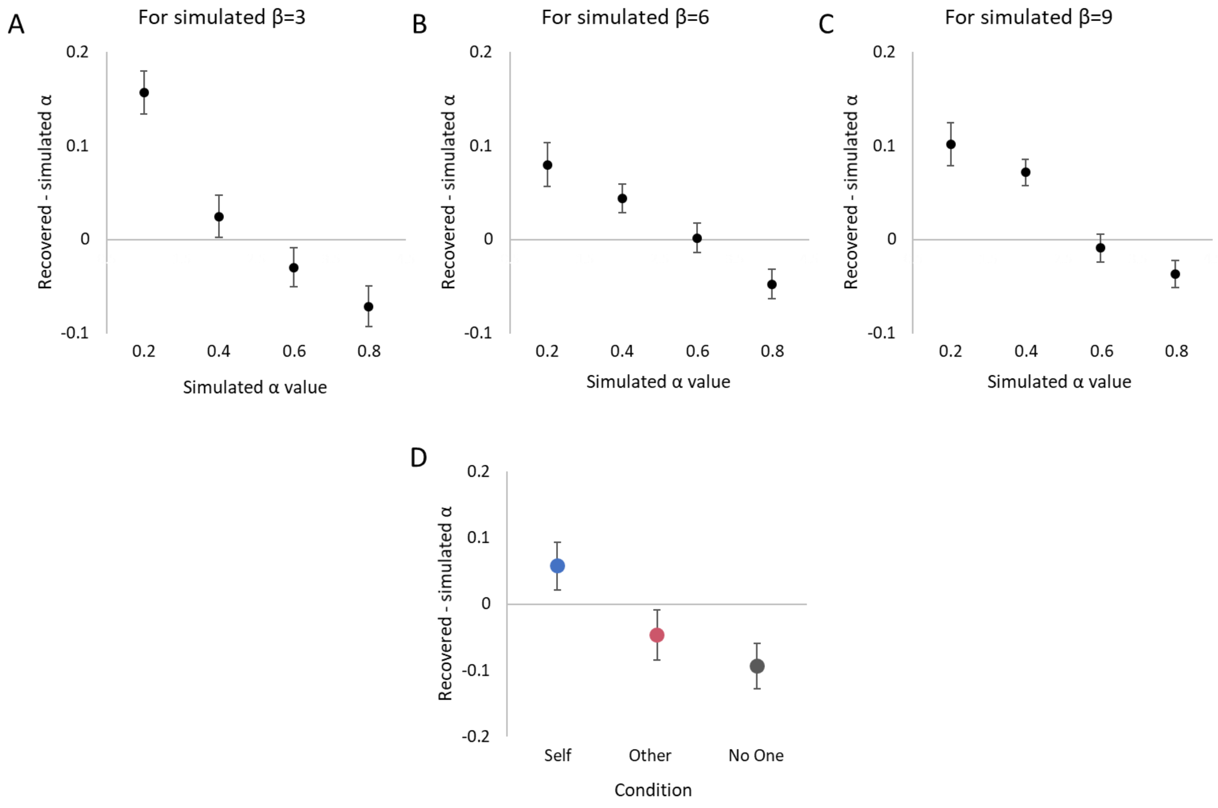


**Figure S4. Learning rate parameter recovery.** The upper panel **(A-C)** shows the recovery of the different learning rates values in models with different β values (i.e., 3, 6, or 9). **(D)** Parameter recovery from a simulation based on real participant data. In all plots, the y-axis indicates how much the recovered values deviate from the true simulated values; the closer to zero the better the recovery.

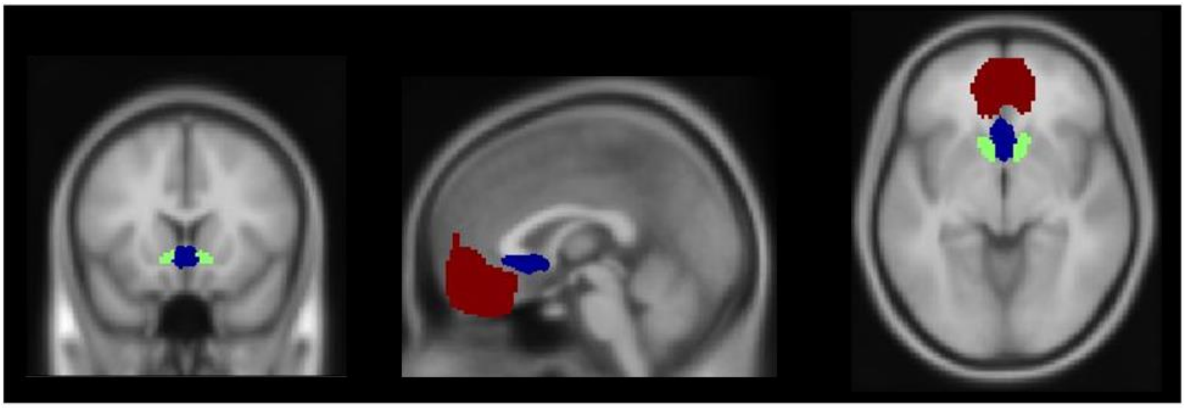

**Figure S5. Regions of interest.** Ventromedial prefrontal cortex (vmPFC; red), subgenual anterior cingulate cortex (sgACC; blue), and the ventral striatum (green).

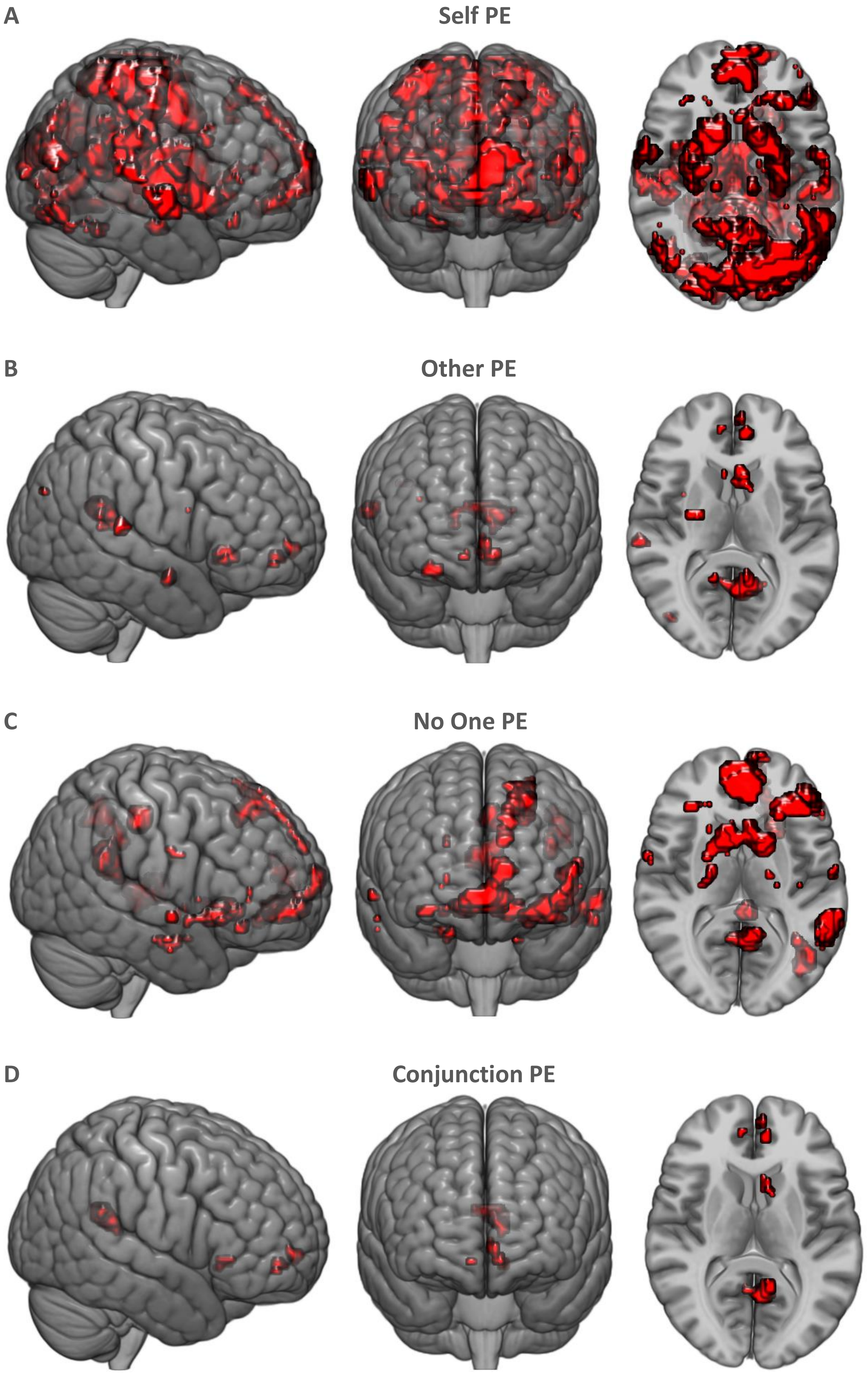


**Figure S6.** Whole brain responses for (A) Self PE, (B) Other PE, (C) No One PE, and (D) conjunction (common PE coding in all three conditions). All images displayed at $p < .05$ FWE, voxel level corrected.

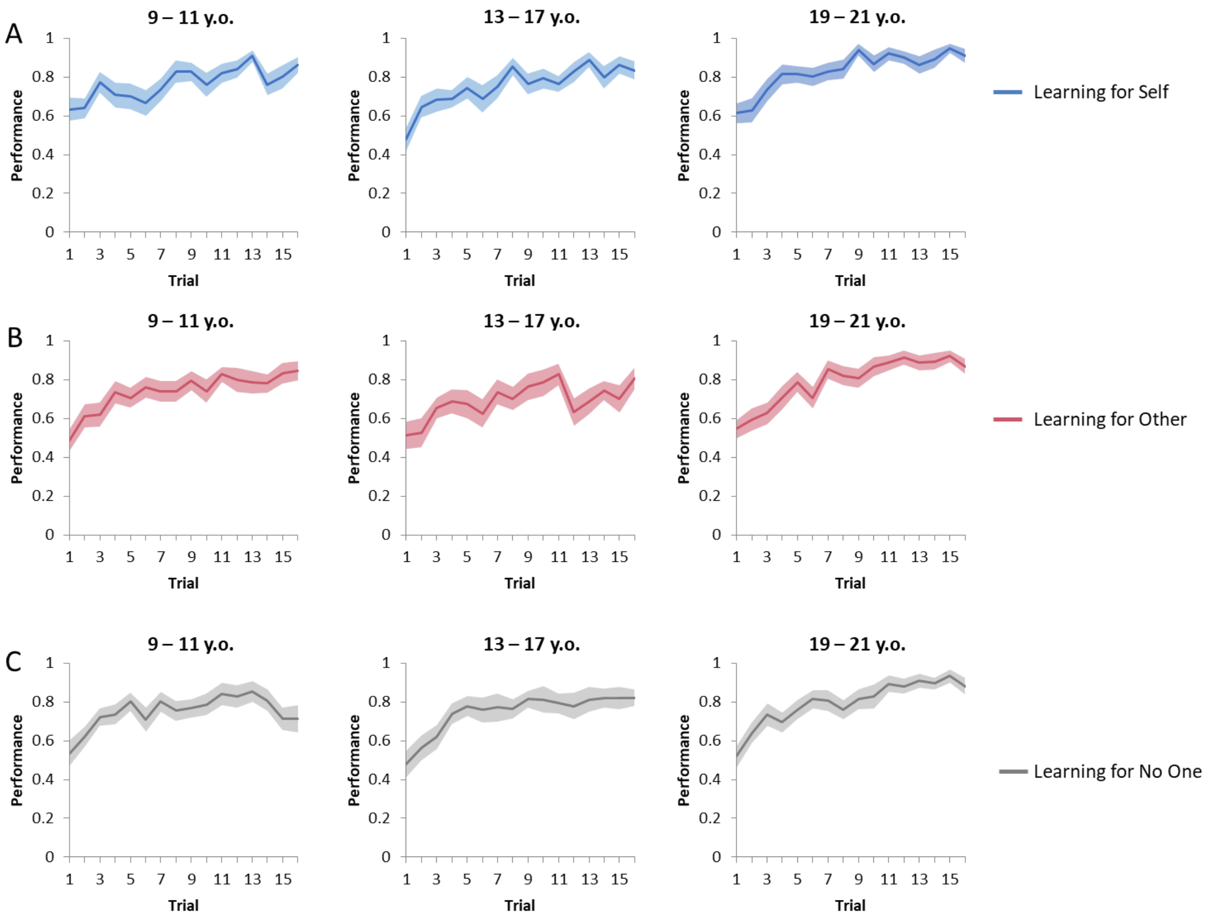


**Figure S7.** Learning across trials for **(A)** Self, **(B)** Others, and **(C)** No One, per age cohort. Note that for all analyses including age, we used age as continuous variables. However, figures represent age per age cohorts instead for illustrative purposes and interpretability.

**Table S1.** Main effects of whole brain prediction error responses per condition, and common prediction error coding (conjunction).

| Brain Region | Peak voxel *x* | *y* | *z* | *k* | *t* | *z* |
|---|---|---|---|---|---|---|
| **Main effect Self PE** | | | | | | |
| L Precental gyrus | -27 | -25 | 61 | 6235 | 9.68 | Inf |
| L precuneus | -3 | -58 | 13 | | 9.65 | Inf |
| L Postcentral gyrus | -30 | -34 | 64 | | 9.03 | Inf |
| L Putamen | -30 | -13 | 4 | 481 | 9.52 | Inf |
| L Putamen | -15 | 8 | -11 | | 8.20 | 7.65 |
| R Caudate | 12 | 11 | -11 | 525 | 9.01 | Inf |
| R Thalamus | 30 | -16 | 7 | | 8.22 | 7.65 |
| R Putamen | 27 | -7 | 10 | | 8.04 | 7.52 |
| L Superior frontal gyrus, medial | -9 | 65 | 19 | 515 | 7.25 | 6.85 |
| L Superior frontal gyrus, medial | -12 | 58 | 7 | | 6.93 | 6.58 |
| L Superior frontal gyrus, medial | 0 | 59 | 1 | | 6.85 | 6.52 |
| L Rolandic operculum | -48 | -28 | 19 | 271 | 6.95 | 6.60 |
| L Superior temporal gyrus | -60 | -31 | 19 | | 6.71 | 6.39 |
| L Supramarginal gyrus | -60 | -22 | 16 | | 6.57 | 6.27 |
| L Middle frontal gyrus, orbital part | -30 | 35 | -14 | 50 | 6.73 | 6.41 |
| L Thalamus | -12 | -22 | 4 | 21 | 6.57 | 6.27 |
| R Superior temporal gyrus | 63 | -1 | -5 | 83 | 6.52 | 6.23 |
| R Rolandic operculum | 63 | 5 | 1 | | 6.06 | 5.82 |
| R Superior temporal gyrus | 66 | -7 | 4 | | 5.29 | 5.13 |
| L Inferior frontal gyrus,triangular part | -48 | 32 | 13 | 60 | 6.47 | 6.19 |
| L Rolandic operculum | -54 | -4 | 4 | 34 | 6.04 | 5.81 |
| R Inferior temporal gyrus | 51 | -67 | -11 | 46 | 5.76 | 5.55 |
| R Inferior temporal gyrus | 51 | -58 | -20 | | 5.41 | 5.23 |
| R Inferior occipital gyrus | 42 | -76 | -17 | | 5.35 | 5.19 |
| R Cerebellum | 21 | -52 | -23 | 20 | 5.60 | 5.41 |
| **Main effect Other PE** | | | | | | |
| L Precuneus | -6 | -61 | 13 | 103 | 6.54 | 6.25 |
| L Calcarine fissure & surrounding cortex | -12 | -55 | 10 | | 6.19 | 5.94 |
| L Olfactory cortex | -6 | 20 | -11 | 32 | 6.29 | 6.02 |
| R Hippocampus | 30 | -7 | -20 | 17 | 5.66 | 5.46 |
| R Superior temporal gyrus | 63 | -28 | 16 | 17 | 5.59 | 5.40 |
| L Middle frontal gyrus, orbital part | -9 | 44 | -11 | 12 | 5.58 | 5.39 |
| **Main effect No One PE** | | | | | | |
| L Middle frontal gyrus, orbital part | -3 | 50 | -11 | 246 | 9.13 | Inf |
| L Superior frontal gyrus, medial | -6 | 62 | 1 | | 5.92 | 5.69 |
| L Superior frontal gyrus, medial | -9 | 55 | 13 | | 5.77 | 5.56 |
| R Caudate | 12 | 8 | -11 | 173 | 7.64 | 7.19 |
| L Olfactory cortex | -15 | 11 | -14 | | 6.72 | 6.40 |
| L Olfactory cortex | -5 | 20 | -11 | | 6.19 | 5.93 |
| L Inferior frontal gyrus, orbital part | -36 | 35 | -14 | 182 | 7.60 | 7.15 |
| L Middle frontal gyrus, orbital part | -24 | 32 | -17 | | 6.30 | 6.04 |
| L Inferior frontal gyrus, triangular part | -45 | 32 | 7 | | 5.99 | 5.75 |
| L Middle temporal gyrus | -60 | -43 | -8 | 134 | 7.21 | 6.82 |
| L Inferior temporal gyrus | -54 | -52 | -17 | | 6.39 | 6.09 |

| | | | | | | |
|---|---|---|---|---|---|---|
| L Precuneus | -6 | -55 | 16 | 120 | 7.16 | 6.78 |
| L Calcarine fissure & surrounding cortex | -12 | -52 | 7 | | 5.88 | 5.66 |
| L Median cingulate and paracingulate gyri | -3 | -37 | 40 | 50 | 6.57 | 6.27 |
| L Middle frontal gyrus | -24 | 32 | 49 | 172 | 6.56 | 6.26 |
| L Middle frontal gyrus | -24 | 20 | 46 | | 5.99 | 5.75 |
| Superior frontal gyrus, medial | -12 | 59 | 28 | | 5.64 | 5.45 |
| L Angular gyrus | -42 | -67 | 34 | 122 | 6.14 | 5.89 |
| R Parahippocampal gyrus | 18 | -10 | -26 | 16 | 5.63 | 5.43 |
| R Parahippocampal gyrus | 24 | -19 | -23 | | 5.20 | 5.04 |
| **Conjunction** | | | | | | |
| L Precuneus | -6 | -58 | 16 | 61 | 6.54 | 6.24 |
| L Caudate | -6 | 14 | -8 | 12 | 5.67 | 5.49 |

For all regions, FWE $p < .05$ voxel-level whole-brain corrected, and presented here with $k > 10$. PE = Prediction error; L = Left; R = Right; k =cluster extent. Names of the brain regions derived from the Automated Anatomical Labeling (AAL) atlas.

**Table S2.** Comparison of responses to prediction errors between conditions, in regions of interest (ventral striatum, sgACC, vmPFC).

| | *Peak voxel* | | | | | |
|---|---|---|---|---|---|---|
| *Brain Region* | *x* | *y* | *z* | *k* | t | z |
| **No One PE > Other PE** | | | | | | |
| Ventral striatum | 12 | 8 | -11 | 2 | 3.41 | 3.36 |
| **No One PE > Self PE** | | | | | | |
| No suprathreshold voxels | | | | | | |
| **Other PE > Self PE + No One PE** | | | | | | |
| No suprathreshold voxels | | | | | | |
| **Self PE + No One PE > Other PE** | | | | | | |
| Ventral striatum | 12 | 8 | -11 | 8 | 4.52 | 4.42 |
| sgACC | 9 | 8 | -11 | 2 | 3.75 | 3.69 |
| **Self PE > Other PE + No One PE** | | | | | | |
| Ventral striatum | 12 | 11 | -11 | 3 | 3.34 | 3.30 |
| **No One PE > Self PE + Other PE** | | | | | | |
| No suprathreshold voxels | | | | | | |
| **Self PE + Other PE > No One PE** | | | | | | |
| No suprathreshold voxels | | | | | | |
| **Other PE + No One PE > Self PE** | | | | | | |
| No suprathreshold voxels | | | | | | |

For all regions, corrected at $p < .05$ FWE-SVC. PE = Prediction error; k =cluster extent. Names of the brain regions were based on the Automated Anatomical Labeling (AAL) atlas.

### Participant instructions

"Welcome! We are going to play a game in the scanner. In this game, you will see two pictures on the screen. You can win or lose points by choosing one of the pictures. If you win, you get +1 point, and if you lose, you get -1 point. But not all pictures are equally good…"

"With both pictures you can win and lose, but with one picture you will win more often, and with the other picture you will lose more often. Try to win as many points as possible! Note: it does not matter whether the picture is on the left or right side of the screen."

"To choose the left picture, you press the left button. To choose the right picture, you press the right button. At the end of the game, you will see how many points you won in total. Your points will be translated to real money using a formula. This amount of money will be paid out to you."

"You will play this game 3 times: for yourself, for another person, and for no one. On the screen, it says for whom you will be playing. Each time you should learn which of the two pictures on the screen is better. Sometimes you play for yourself. When you play for yourself, the gains will be paid out to you."

"Sometimes you play for another person. When you play for another person, the gains will be paid out to another player. This player is someone who participates in this experiment after you. This is a girl or a boy of your age. This person does not know that you are playing for him/her. So, he/she will receive the money you win for him/her without them knowing it is from you. This person will not play the game for you."

"Sometimes you play for no one. When you play for no one, your points don't count and no one will receive your gains."

"Try to respond on time. You will have about 2 seconds to make your choice. We will first do a practice run. Good luck!"

*[24 Practice trials (8 per condition)]*

"Well done! This was a practice run, so your points don't count yet. In the scanner, we will play the game for real, and you will see at the end of the game how many points you won for yourself and for the other person. Do you have any questions left?"
